# Comprehensive Structural Variant Detection: From Mosaic to Population-Level

**DOI:** 10.1101/2022.04.04.487055

**Authors:** Moritz Smolka, Luis F. Paulin, Christopher M. Grochowski, Dominic W. Horner, Medhat Mahmoud, Sairam Behera, Ester Kalef-Ezra, Mira Gandhi, Karl Hong, Davut Pehlivan, Sonja W. Scholz, Claudia M.B. Carvalho, Christos Proukakis, Fritz J Sedlazeck

## Abstract

Long-read Structural Variation (SV) calling remains a challenging but highly accurate way to identify complex genomic alterations. Here, we present Sniffles2, which is faster and more accurate than state-of-the-art SV caller across different coverages, sequencing technologies, and SV types. Furthermore, Sniffles2 solves the problem of family- to population-level SV calling to produce fully genotyped VCF files by introducing a gVCF file concept. Across 11 probands, we accurately identified causative SVs around *MECP2*, including highly complex alleles with three overlapping SVs. Sniffles2 also enables the detection of mosaic SVs in bulk long-read data. As a result, we successfully identified multiple mosaic SVs across a multiple system atrophy patient brain. The identified SV showed a remarkable diversity within the cingulate cortex, impacting both genes involved in neuron function and repetitive elements. In summary, we demonstrate the utility and versatility of Sniffles2 to identify SVs from the mosaic to population levels.

## Introduction

The role and biological impact of Structural Variations (SVs) have become evident^1,2^. SVs are loosely defined as 50 bp or larger genomic alterations that fall into five types (insertions, inversions, deletions, duplications, and translocations) or a combination of these types^1^. Given that this type of variant impacts the most number of nucleotides in a genome, it is not surprising that evidence is mounting regarding their importance across all categories of life. This starts e.g., with important speciation events^3^ and impacts plants^4,5^, but goes further across human diseases (Mendelian^6,7^ as well as complex diseases^8–10^) to cancer development^11–13^ (e.g., HLA loss, oncogene amplification). Despite the importance of SVs, it is still challenging to detect germline and somatic SVs or even robustly identify *de novo* SVs^14–16^. The least often studied and thus challenging SVs are insertions (ie. novel sequences) that, as many studies showed, amount to half of all SVs found in a human genome^17–19^. The latter can either be recovered by long-read mapping methods or *de novo* assemblies followed by a genomic alignment^1,20^.

Long-read sequencing has come a long way over the past years from a novelty to a population/production scale mechanism to study SVs^21,22^. The error rate of Oxford Nanopore and PacBio HiFi are both ever decreasing, soon reaching levels of Illumina-like errors along the genome^23,24^. Most recently, Oxford Nanopore Technologies (ONT) provided an insight into the upcoming chemistry update to produce Q20+ reads, which further seem to reduce the error rates (∼2%)^25^. Indeed, several studies have now started to sequence larger and larger data sets or even medical applications using PacBio HiFi or ONT^21,26^. This trend started with GENCODE ^22^, but is ever increasing to other projects (e.g., All of Us initiative^27^, CARD^28^) and is currently peaking in the G42 endeavor to sequence multiple hundreds of thousands genomes. This also requires more efficient software to not just detect SVs, but also to merge and produce a fully genotyped VCF file^29,30^. The improved degrees of error and lower cost for long-reads are also starting to promote applications in medical or clinical space^31,32^. This is needed as several genes or regions of the genome remain a “dark matter”^20,33^. Here, recent studies showed 386 medically relevant genes that are still challenging for the standard clinical Illumina WGS^33^. Most of these genes (∼70%) can be assessed using long-read technologies, but several challenges remain^33^.

Furthermore, there are more complex SVs beyond simple deletions, duplications, inversions, insertions, and translocations that can lead to a Mendelian disease^6^. The genomic locus including the dosage-sensitive gene *MECP2* at Xq28 is particularly susceptible to such genomic instability due to nearby inverted and direct orientation low-copy repeats (LCRs)^34–36^. The protein encoded by the *MECP2* gene, Methyl-CpG binding protein 2 (MeCP2), is critical for brain function by acting as an epigenetic regulator^37^. Copy-number variation spanning the gene causes *MECP2* Duplication Syndrome (MDS) (MIM:300260) with 100% penetrance in males^38^. The most prevalent clinical features of MDS are infantile hypotonia, developmental delay, intellectual disability, frequent respiratory infections, and refractory epilepsy^39^. One of the frequent complex allele presentations is constituted by an inverted triplication flanked by duplications (DUP-TRP/INV-DUP). This allele is generated by a given pair of inverted low-copy repeats telomeric to *MECP2*, being responsible for 20% to 30% of the MDS cases^6^ a fraction of which will lead to a more severe clinical phenotype. When generated, this structure includes two breakpoint junctions (Jct) connecting the end of the duplication to the end of the triplication (Jct1) and the beginning of the triplication to the beginning of the duplication (Jct2). Given the presence of two breakpoint junctions in cis, the involvement of LCRs and the size of such events (often > 500 kb), we lack the ability not only to detect this structure solely using long read sequencing data, but also how to describe it following the VCF specification. Part of the complexity originates as the reads themselves only partially indicate the allele, e.g., highlighting a shorter inversion^29^.

In addition to complex variants, multiple studies have shown that there are mosaic or low-frequency SVs that are likely causal across neurological diseases or other diseases^9^. As an example, single-cell studies show us that there can be variable copy number variants (CNVs) across multiple cells in the brain^9^. However, their true frequency is unknown, with around 12% of healthy cortical neurons having Mb-scale CNVs^40^. A possible role in neurodegenerative disease^41^ has not been adequately explored. In synucleinopathies, which include Parkinson’s disease and Multiple System Atrophy^42^ (MSA), somatic CNVs of the highly relevant *SNCA* gene have been reported^43–45^, and single-cell whole-genome sequencing in MSA has shown Mb-scale CNVs in ∼30% of cells^44^. Still, these CNVs studies lack resolution as breakpoints are defined within +/- multiple kbp and only very large ∼1 Mb CNV events are reported^40,46,47^. An identification of complex SVs arising in neurodevelopment was so far only possible with WGS of clonally-expanded precursors^9,44^. Thus, so far we struggle to identify the underlying alleles even for large already reported CNVs along the human genome.

In this paper, we present Sniffles2, a redesign of the popular SV caller Sniffles, with improved accuracy, higher speed and novel features that addresses the problem of population-scale SV calling for long-reads. This is needed across tumor-normal comparison over family (e.g. mendelian) studies, but also at larger studies deciphering rare alleles across a population or cohort. In addition, Sniffles2 enables the detection of low-frequency SVs across data sets, which facilitates detection of somatic SVs and mosaicism studies and opens the field of cell heterogeneity for long-read applications. We first highlight Sniffles2 performance compared to other SV caller over multiple benchmark sets then we further investigate how the new population or family mode for SV calling improves the accuracy and performance across Mendelian disease probands with ONT. Here, we can showcase the boundaries of long-read SV calling by assessing highly complex SVs around *MECP2.* Lastly, we investigate Sniffles2 abilities to identify low-frequency/mosaic SV across an MSA brain sample and compare its performance to Illumina sequencing and Bionano optical genome mapping (OGM). Overall, Sniffles2 pushes the boundaries of long-read based SV calling and thus demonstrates the utility of such an approach further than any existing approach. Sniffles2 remains an open source (MIT license) and is available at: https://github.com/fritzsedlazeck/Sniffles

## Results

### Accurate detection of structural variations at scale

Sniffles2 is a complete redesign and extension of the popular SV caller Sniffles. **Figure 1** gives an overview of its main components. Sniffles2 now implements repeat aware clustering to improve germline SV calling (**Figure 1A**) and further enables family and population SV calling at scale and ease (**Figure 1B**) and implements novel methods to identify mosaic SVs (**Figure 1C**). A detailed description of Sniffles2 can be found in the methods section.

**Figure 1:**
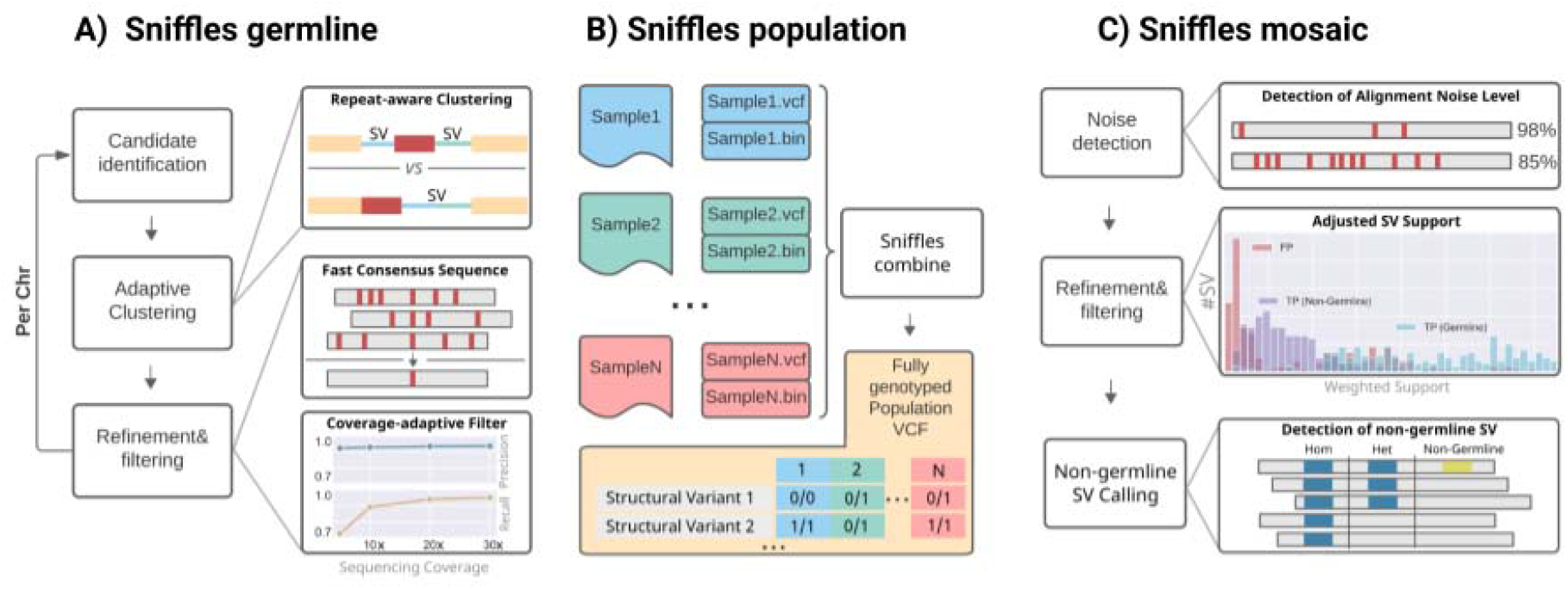
Overview of Sniffles2. **A)** For Sniffles 2 we have implemented a repeat aware clustering coupled with a fast consensus sequence and coverage-adaptive filtering to improve accuracy of the germline SV calls. **B)** One key limitation of current SV calling is the generation of fully genotyped population VCF. Sniffles2 implements a concept similar to a gVCF file where single sample calling is only done once which reduces runtime multiple-fold. **C)** Mosaic SV detection is enabled by improved detection and filtering of low variant allele frequency SV (by default 5-20%) across a bulk sample. This is enabled over additional noise detection methodology as well as refinement and filtering approaches that we developed.

**Figure 1A** shows a summary of the most important steps applied by Sniffles2 to identify germline SVs. In brief, we use a fast yet high-resolution clustering approach, which identifies SVs in three key steps. First, putative SV events are extracted from read alignments (split reads and inline insertion or deletion events) and allocated to high-resolution bins (default: 100 bp) based on their genomic coordinates and putative SV type. Second, neighboring SV candidate bins are subsequently merged based on a standard deviation measure of SV starting positions within each growing bin. Through the use of optional tandem repeat annotations, Sniffles2 dynamically adapts clustering parameters during SV calling, allowing it to detect single SVs that have been scattered as a result of alignment artifacts. Finally, identified clusters are separately reanalyzed and split based on putative SV length. Final SV candidates are subjected to quality control based on read support, breakpoint variance and expected coverage changes.

We assessed the performance of Sniffles2 (v2.2) with respect to Sniffles^29^ (v1.12), cuteSV^48^ (v1.0.11), PBSV^49^ (v2.6.2) and SVIM^50^ (v1.4.2) using Truvari^51^ (version 2.1) and the GIAB recommended parameters^52^. **Figure2** shows the results across different GIAB benchmarks see **Supplementary Table 1** for details). Across the default coverage (30x for HiFi, 50x for ONT), Sniffles2 shows the best performance with respect to correctly identified and genotyped insertions (HiFi: GT F-score 0.909, ONT: GT F-score 0.915) and deletions (HiFi: GT F-score 0.934, ONT: GT F-score 0.944) (see **Supplementary Table 2** for details). Sniffles2 achieves a better result in a fraction of the time across data sets compared to Sniffles (v1.12), being over 17 times (HiFi) and 11 times (ONT) faster in processing a 30x coverage data set, respectively. Figure 2A**+B** shows the results for PacBio HiFi and ONT, respectively. In addition, Sniffles2 is also the fastest method overall, requiring 33.33 CPU minutes for processing a 30x coverage HiFi dataset , which was twice as fast as SVIM, the 2nd fastest method. For a 30x coverage ONT dataset, Sniffles2 was also close to twice as fast (1.92x) as the second fastest caller (SVIM), while also having an over 8.4% higher GT F-score. Taking into account Sniffles2 multi- processing capability (not supported by SVIM), the speedup increases even further, to more than 5.4-fold and 7.5-fold for HiFi 30x, ONT 30x data sets, respectively. When reducing the coverage from 30x to 10x we observe only a slight reduction in F-score for Sniffles2 (HiFi: reduction GT F-score 0.041, ONT: reduction F-score 0.058). This is in stark contrast to other programs such as cuteSV, where using default parameters, F-score dropped by an average of over 60% (HiFi: reduction F-score 0.56, ONT: reduction F-score 0.58). Even when using only 5x for Sniffles2, we still observe a high accuracy ONT (F-score: 0.74) and HiFi (F-score: 0.77). This is achieved as Sniffles2 includes an automated parameter selection for filtering of SV candidates based on the available coverage. In contrast, other SV callers rely on manual adjustment of these parameters to retrieve acceptable results across coverages and sequencing technologies. Figure 2A-D shows this clearly as all other SV callers show a decreased performance across lower coverage. Even when tuning the parameters for other SV callers (Figure 2A-D, Tuned parameters), Sniffles2 remains the highest accuracy (see **Supplementary Table 1** for details).

**Figure 2:**
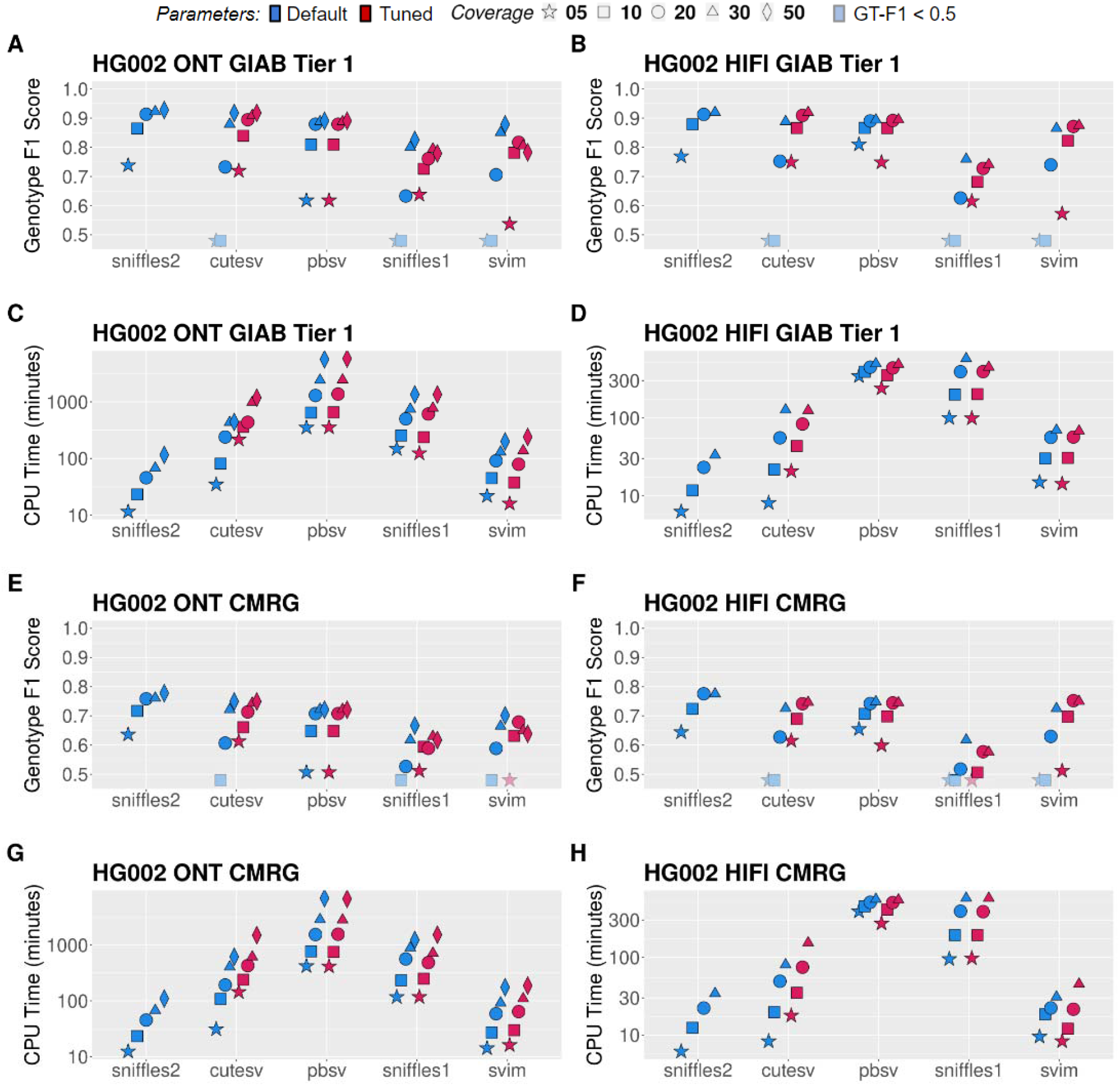
Performance assessment of Sniffles2 based on GIAB. Performance metrics for correctly identifying and genotyping SVs across Oxford Nanopore (left) and PacBio HiFi (right). All details are collected in **Supplementary Table 1**. For panels A, B, E and F the shaded symbols mean that the Genotype F1 score was lower than 0.5. **A+B:** comparison across Tier1 GIAB genome-wide SV (Genotype F1 score in the y-axis, higher is better) across different coverages (symbols) and SV caller (x-axis) for default and maximum sensitivity parameters (red and green respectively / clear and solid respectively). **C+D:** runtime comparison across Tier1 GIAB genome-wide SV (CPU minutes in the y-axis, lower is better) across different coverages (symbols) and SV caller (x-axis) for default and maximum sensitivity parameters (red and green respectively / clear and solid respectively). **E+F:** comparison across GIAB challenging medical gene (CMRG) benchmark for SV (Genotype F1 score in the y-axis, higher is better) across different coverages (symbols) and SV caller (x-axis) for maximum sensitivity parameters. **G+H:** runtime comparison across GIAB CRMG benchmark for SV (CPU minutes in the y-axis, lower is better) across different coverages (symbols) and SV caller (x-axis) for maximum sensitivity parameters.

Additionally, we tested Sniffles2 with an additional mapper (LRA) to showcase its versatility. When mapping HiFi data with LRA we observe a slight increase in performance when comparing the genotype F-scores (HiFi GT F-Score increase by 0.006), moreover when using LRA to map ONT data we observe a slight decrease in performance of Sniffles2 (ONT GT F-Score decrease 0.0136). Interestingly, when testing Sniffles with CLR data the error rates of this outdated data impacts our variant calling and thus we suggest other SV callers (see **Supplementary Table 1**).

Next, we performed an evaluation with respect to the Tier2 GIAB dataset, which is a more challenging region of the GIAB benchmark set as it includes repeats and GIAB cannot guarantee the accuracy of the variants in these regions (see **Supplementary Table 3** for details). Again, Sniffles2 even increases the performance difference compared to other SV caller. Lastly, we benchmarked Sniffles2 across a more challenging SV data set across 386 medically relevant, but highly polymorphic/challenging genes^33^. GIAB has recently released this call set of ∼200 SV covering around 70% of these genes^33^. Figure 2E-H shows the results. Again Sniffles2 outperforms the other SV callers in terms of accuracy and speed using default parameters. The next best performing SV caller (pbsv for HiFi, cuteSV for ONT) both achieved 3.6% and 5.5% lower genotyping accuracy (GT F1-Score) even at 30x coverage. **Supplementary Table 4** contains the detailed results across all SV callers. Overall, Sniffles2 outperforms other state-of-the-art SV caller across the entire genome including the most challenging regions/genes.

Next, Sniffles2 improves insertion identification through two additional methods: First, the consensus module corrects sequencing-related errors in the recovered insertion sequences using a fast pseudo-alignment based approach. This allows Sniffles2 to attain the second highest mean sequence identity of (HiFi: 0.948, ONT: 0.939), after pbsv (HiFi: 0.953, ONT: 0.949), while Sniffles2 is over 14x (HiFi) and 36x (ONT) faster (see **Supplementary** Figure 1**, Supplementary Table 5** containing insertion sequence accuracy across all callers) at 30x coverage. Second, Sniffles2 increases the sensitivity for the detection of large insertions by recording additional supporting alignment signals in the affected regions (see **Supplementary** Figure 2**, Supplementary Table 6**) at much higher speed than pbsv, the only SV caller with a comparable accuracy for long insertions.

Lastly, GIAB only represents one individual benchmarked across most studies (HG002). Thus next, we used Dipcall^53^ together with three T2T assemblies ^54^ (HG01243, HG02055, HG02080 of America, African and East-asian ancestry respectively) to further assess the performance of Sniffles2. Clearly, we give Dipcall the benefit of the doubt, knowing that the accuracy will be lower than the GIAB vetted benchmark set. Overall, Sniffles2 performs the best across all samples having on average a F-Score of 0.79 at 30x coverage ONT and HiFi compared to 0.77 F-score for the next best SVcaller (cuteSV), at nearly 3.5 times the speed (74.03 CPU minutes versus 256.65 CPU minutes on average) for default parameters. **Supplementary Table 7** contains the detailed results for each benchmarked program across these three samples. Besides the here benchmarked insertions and deletions, we also benchmarked Sniffles2 on duplications, inversions and translocations using simulated data as no benchmark exists. Overall, Sniffles2 again outperformed all other methods in speed and accuracy (see **Supplementary** Figure 3 **and Supplementary Table 8**) (see methods for details).

Given all these comparisons across different ethnicities (HG002 being European, HG01243 being American, HG02055 being African and HG02080 being East-Asian), coverage levels (5- 50x) and sequencing technologies (HIFI and ONT), we conclude that Sniffles2 improves the detection of SVs in terms of accuracy and speed compared to other state-of-the-art methods.

### Enabling family to large cohort studies to discover the impact of complex Structural Variation

Over the past years, an uptake of ever larger studies utilizing long-reads is foreshadowing a trend in genomics to utilize long-reads more often than ever^21^. To promote this, Sniffles2 is fast and efficient, but further implements a strategy to obtain a fully genotyped population VCF. Traditionally this is a multi-stage process of calling, merging, genotyping, and re-merging^21,55,56^. This is clearly inefficient as the bam/cram alignment files need to be assessed twice. Even so, this process can only be achieved by using a few of the existing methods (SVJedi^57^, Sniffles^29^, CuteSV^48^). Sniffles2 strategy only requires an initial calling and merging to obtain a fully genotyped population-level VCF. Figure 1B illustrates the principle. The calling can be done independently per sample and thus allows to scale to large data sets. Each sample run produces a single germline VCF file accompanied with a binary file that serializes every single candidate SV that has even a single read support. Next, each binary file per sample is provided as a list to Sniffles2 merge, which combines the SV across the samples and fills the missing information utilizing the binary files per sample. These files typically range from 75 to 250Mb per ∼30x ONT sample. This process is extremely efficient as it scales linearly with the number of samples and allows the samples to be analyzed in parallel and independent of each other (see **Supplementary table 9, Supplementary** Figure 4). In addition, it solves the “n+1” problem to include a batch of samples at a later stage of a project.

To assess the performance of this approach, we measured the Mendelian inconsistency rate using a family trio (see methods)^58^. Here we counted in how many of the called SV, the genotypes of the proband do not concord with the genotype of the parents (e.g., F 0/0, M 0/0, P 1/1) or the other way around (e.g., F 1/1, M 0/0, P 0/0). For Sniffles2, we obtained a Mendelian inconsistency rate of 9.13% with a low rate of missing genotype of 1.29% (Figure 3A). The latter is driven by a user parameter where Sniffles2 does not report a genotype where only 5 or fewer reads are present. In comparison, cuteSV with a simple merge (SURVIVOR^59^) presented a mendelian inconsistency of 3.74% with a much higher missingness of 32.20% of all genotypes compared to the 1.29% of Sniffles2. When we apply a re-genotyping and re-merging of the cuteSV results, we obtain a Mendelian inconsistency rate of 8.88% with almost 3 times higher missingness of 3.45%, when compared to Sniffles2. Furthermore, the cuteSV approach took more than 50 hours CPU hours (50:33:13, **Supplementary Table 10, Supplementary** Figure 5) in contrast to 8 CPU hours (8:05:34) for Sniffles2 for a given trio. Thus, rendering it impractical for larger cohorts. As a stress test we merged from 3 to 777 samples which consisted of repeating up to 259 times the HG002 family trio. This took little more than 11 CPU hours using Sniffles2 (11:16:18, **Supplementary table 9**).

**Figure 3:**
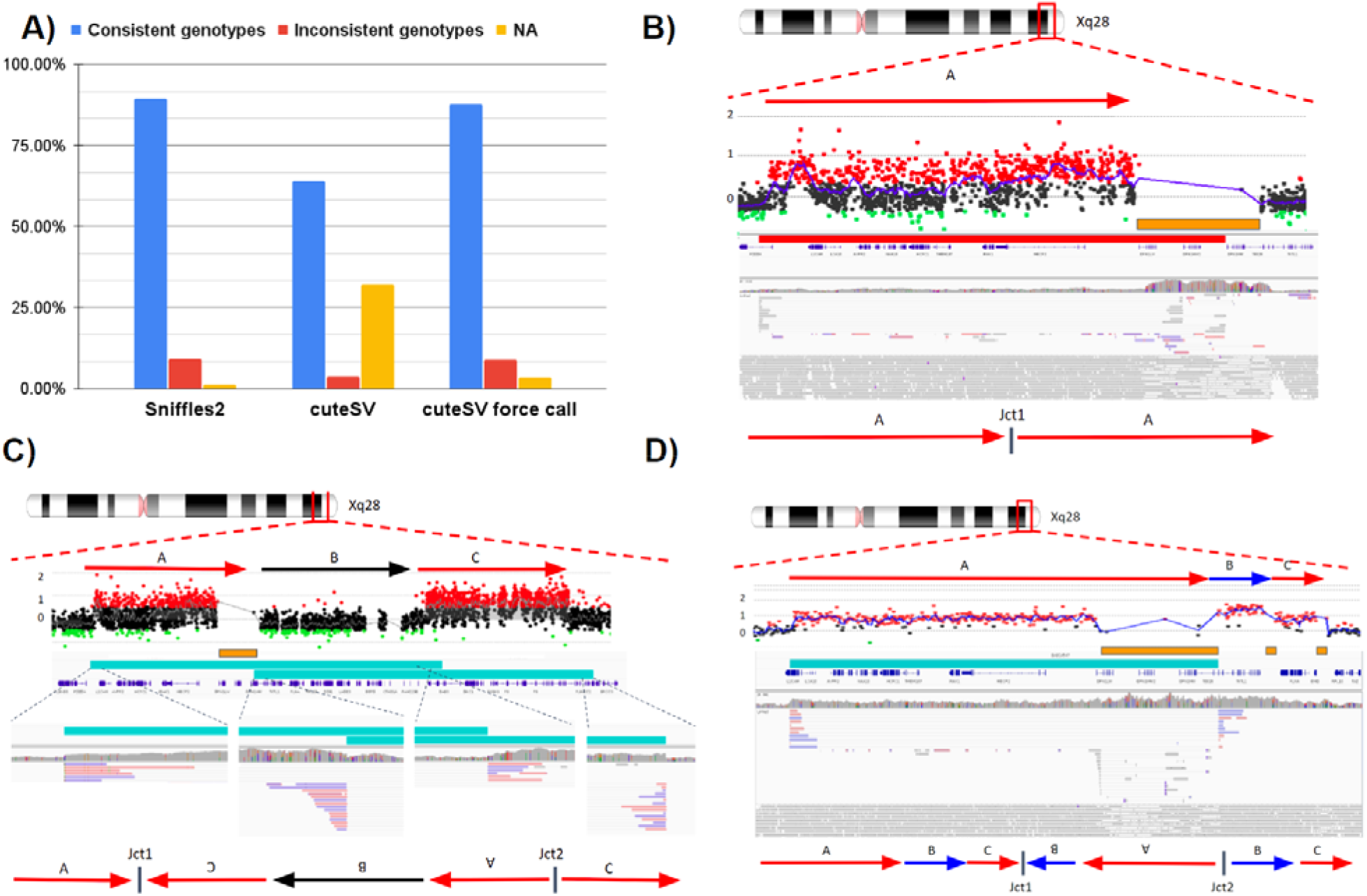
Sniffles2 population approach and application to Mendelian disease. **A** Comparison of the proportion of consistent, inconsistent and uninformative (NA) genotypes across HG002/3/4 computed by the bcftools Mendelian plugin for Sniffles2 population merge and cuteSV. To achieve similar results, cuteSV takes more than 6.24x the time. **B,C,D)** Three examples of SV detected by Sniffles2 in Mendelian disorders in probands. Chromosomal position is shown in the top part (Xq28), followed by the arrows that represent a specific loci. Next is shown aCGH data dots represent genomic positions being assayed. Black dots represent a log2 ratio between - 0.35 and 0.35, red dots represent a log ratio above 0.35 and green dots represent a ratio below -0.35. Consistent (at least 3 consecutive probes) log2 ratios above 0.35 represent a region of copy number gain and below -0.35 represent copy number loss. In orange we show segmental duplications (SegDups) and in teal the SV called by Sniffles2. IGV screenshot and fully resolved events are shown in the lower part of each example. **B)** Tandem duplication that was fully resolved by Sniffles2 in one of the patients (BH14233_1). Sniffles2 was able to identify and map the junction of the duplication within a segmental duplication region where array data does not provide information. Note that given the presence of the segmental duplication, the SV was tagged with a STDEV_LEN filter. **C)** Detailed aCGH view of a complex duplication-normal-duplication (DUP-NML-DUP) structure in sample BH13947_1 with breakpoints within SegDup or low-copy repeats (LCRs) region (orange bar) where Sniffles2 is indicating two overlapping inversions in IGV (teal bars) forming junctions 1 and 2 (Jct1 and Jct2). Top arrows indicate the reference orientation (duplications in red, neutral regions in black) of each genomic fragment and bottom arrows indicate a possible DUP-NML-INV/DUP haplotype structure containing Jct1 and Jct2. **D)** Sample BH15700_1 shows a complex duplication-triplication-duplication structure as highlighted in aCGH data with SegDups and LCRs highlighted (orange bars). Sniffles2 identifies the inversion breakpoint at Jct2 (teal bar), but cannot fully resolve the entire allele including Jct1 as it’s also not possible to be reported in the VCF standard. Red arrows indicate duplicated regions and blue arrows show triplicated portions. One possible haplotype structure for a DUP-TRP/INV-DUP is shown with the triplication and initial duplication being inverted forming Jct1 and Jct2^35^.

Next, we applied this population/family approach of Sniffles2 across 31 Oxford Nanopore data sets that represented cases of Mendelian disorders in probands (seven complete trios, one duo, eight only probands). The runtime for merging the individual samples was roughly 28 CPU minutes (28:37) to produce a fully genotyped population VCF file. Across the seven complete families (proband, mother, father) we measured an average of 3.89% Mendelian inconsistency rate and 1.11% of missingness (see methods, **Supplementary Table 11, Supplementary** Figure 6). The probands for sequencing were selected based on a Mendelian disease that often is caused by SVs impacting the *MECP2* gene at Xq28 locus. As described in the introduction this is a severe neurodevelopmental disorder that is often caused by extreme complex alleles in this region. We were interested if Sniffles2 together with ONT data can resolve the breakpoints which were not always solvable using array data and if we were able to fully explain the entire allele or just partially solve the junctions. To address this we filtered SV based on ChrX together with their size (10 kb) and filtered for SV only being *de novo* or inherited from the mother.

Within this cohort, Sniffles2 is able to achieve a high rate of detection across junctions, but sometimes struggles to recapitulate the entire allele that contains complex SVs. **Table 1** shows the details per proband. In samples harboring a tandem duplication, Sniffles2 was able to properly detect the allele and fully resolve its architecture. In our cohort, these duplications span the dosage sensitive gene (*MECP2*) and form a single breakpoint junction (Jct1), confirming a tandem duplication structure. As highlighted in sample BH14233_1, although aCGH broadly defines the genomic interval of the duplicated region, Sniffles2 is able to properly give positional context of genomic fragment defining at nucleotide-level resolution to be a tandem duplication on the allele even though the end of the duplication is within a segmental duplication region (orange bar) (Figure 3B). Note that the presence of the segmental duplication caused the SV to be tagged with a STDEV_LEN filter. This indicates non agreement on the precise start of the SV given the repetitive nature of the region.

**Table 1:**
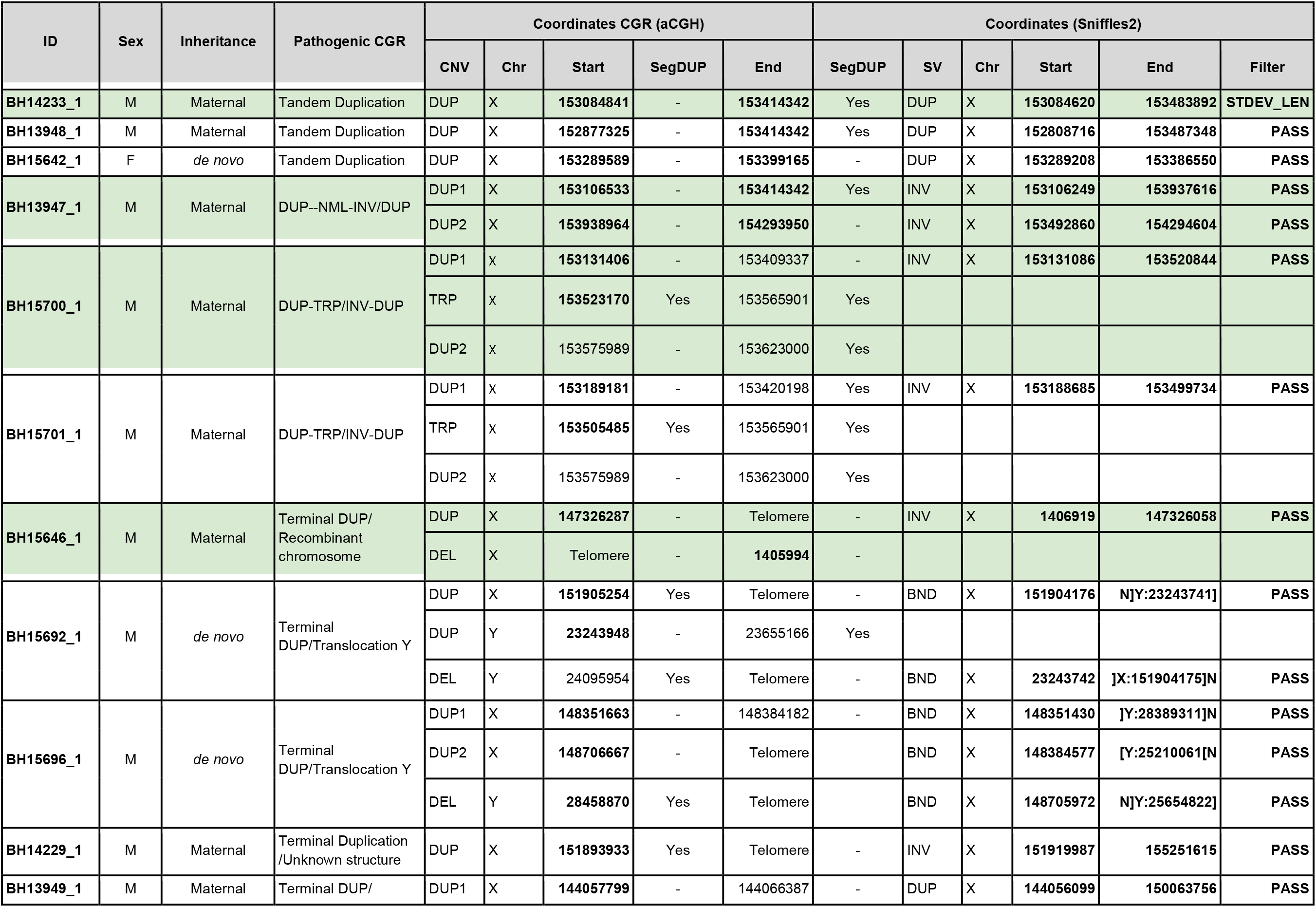
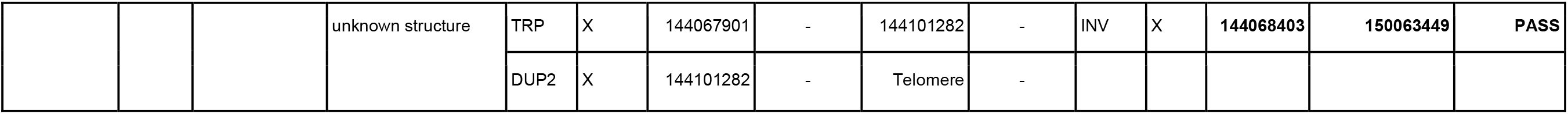
Table across all the probands assessed here and highlighting in bold which junctions could be resolved using Sniffles2. Highlighted in green are the results discussed in the main text.

Interestingly, a portion of the inversions that Sniffles2 was able to detect were not simple genomic inversions, but instead part of more complex structures that could not be fully resolved using current bioinformatic tools. A more complex allele is detected in sample BH13947_1, which consists of a duplication-normal-duplication (DUP-NML-INV/DUP) with breakpoints spanning segmental duplications (SegDups) (Figure 3C). Here Sniffles2 indicates two overlapping inversions which form junctions 1 and 2 (Jct1 and Jct2) generating a DUP-NML-INV/DUP structure.

In sample BH15646_1, the inversion called by Sniffles2 spanning nearly the entire X chromosome (∼148 Mb) represents the breakpoint junction of a recombinant chromosome. In the sample, aCGH data shows a short-arm deletion and a long-arm duplication, i.e. DEL-NML-DUP structure. Sniffles2 is able to positionally connect the beginning of the duplication to the end of the deletion forming Jct1 (**Supplementary** Figure 7). This allele is generated as the result of meiotic recombination between heterozygous homologous X-chromosomes in females harboring a pericentric inversion^60^.

Another example is represented by an apparent 311 kb inversion detected in sample BH15700_1. This inversion is part of a DUP-TRP/INV-DUP structure (Figure 3D), which is generated by a given pair of inverted SegDups and produces an inverted triplication flanked by duplications^35^. When generated, this structure includes two breakpoint junctions (Jct) connecting the end of the duplication to the end of the triplication (Jct1) and the beginning of the triplication to the beginning of the duplication (Jct2). While Sniffles2 can properly detect the inverted breakpoint generating Jct2, it is not able to fully resolve the context of the larger structure due to Jct1 being embedded within a pair of inverted SegDups with 99.9% sequence similarity.

In this cohort, Sniffles2 is able to correctly detect with nucleotide-level resolution the precise breakpoints defining a genomic interval in patients carrying complex genomic rearrangements (CGRs). Importantly, a large portion of the CGRs in this cohort have at least one of the breakpoint junctions mapping to SegDups; those can be fully resolved by Sniffles2 together with copy-number information. Nevertheless, CGRs that have two breaks mapping to pairs of highly identical SegDups, such as in the DUP-TRP/INV-DUP events, still represent an important challenge for SV callers and are also too complex to appropriately report within the limits of a VCF standard. Additionally Sniffles2 infers positional connections that help resolve a given complex allele architecture with information that aCGH alone cannot provide. Thus, overall this highlights the benefit of Sniffles2 family/population mode to enable these types of comparisons.

### Identification of mosaic SVs reveals new insight in diversity

We have shown that Sniffles2 accurately identifies SVs across the entire genome and that it enables better scaling and accuracy across families and even larger population levels. Nevertheless, as we know from many studies, germline variants are not the only source of structural variation. Often somatic/mosaic variants are important. This has been indicated in e.g. cancer and neurological disorders^9,12^. Thus, Sniffles2 is equipped with a mosaic mode to identify low-frequency (5-20% Variant allele frequency (VAF)) SVs across a single sequenced sample. Figure 1C shows the principle steps where the main innovation is to weigh the support of each read taking into consideration its edit distance as a confidence measure. To circumvent the impact of sequencing error rates on mosaic SV detect we filter out SV where the average edit distance of reads supporting exceeds a threshold, which is estimated per data set to account for different sequencing error levels (see methods).

To assess the performance of Sniffles2 across mosaic SVs, we first synthetically merged HG002 (at low concentrations: 5x (7%), 7x (10%), 10x (14%), 15x (21%) or 20x (28%)) with the 50-63x coverage read data from the sample of an unrelated individual (HG00733). This yielded multiple, synthetic samples with constant total coverage of ∼70x (68x in one case), but varying concentrations of HG002 in them (7-28%). We only assessed this for ONT data as this technology often is sequenced at higher sequence depth than Pacbio HiFi. The latter can still be used, but is not benchmarked here. Figure 4 **A&B** highlights the results across this synthetic data set across different concentrations of HG002 (x-axis, **Supplementary Table 12**). Figure 4A **and 4B** show the precision and recall of SVs across the different concentrations. In blue, we highlight the performance of Sniffles2 default germline-mode, which shows low precision and increased recall with increase in concentration of HG002 in the mix. In red we show cuteSV which has lower precision and recall in both cases, when compared to sniffles2 germline-mode. In yellow we highlight the performance of Sniffles2 mosaic mode. Sniffles2 achieves high precision that increases with higher proportions of HG002 (Figure 4A) with the highest being at 21% HG002 proportion (84.12% precision). Recall also increases with higher proportions of HG002, moreover it only reaches a maximum of 54.05%. This is because Sniffles2 mosaic- mode only calls SV where the variant allele frequency (VAF) is between 5 and 20%.. We computed the adjusted recall by only using the SVs where the VAF matches in the range that mosaic-mode uses (Figure 4B**, Supplementary Table 12**). This adjusted recall is shown in green, which averages 94.47% across all comparisons (highest 96.39% at 7% mix of HG002). Thus following the default recall for germline variants.

**Figure 4:**
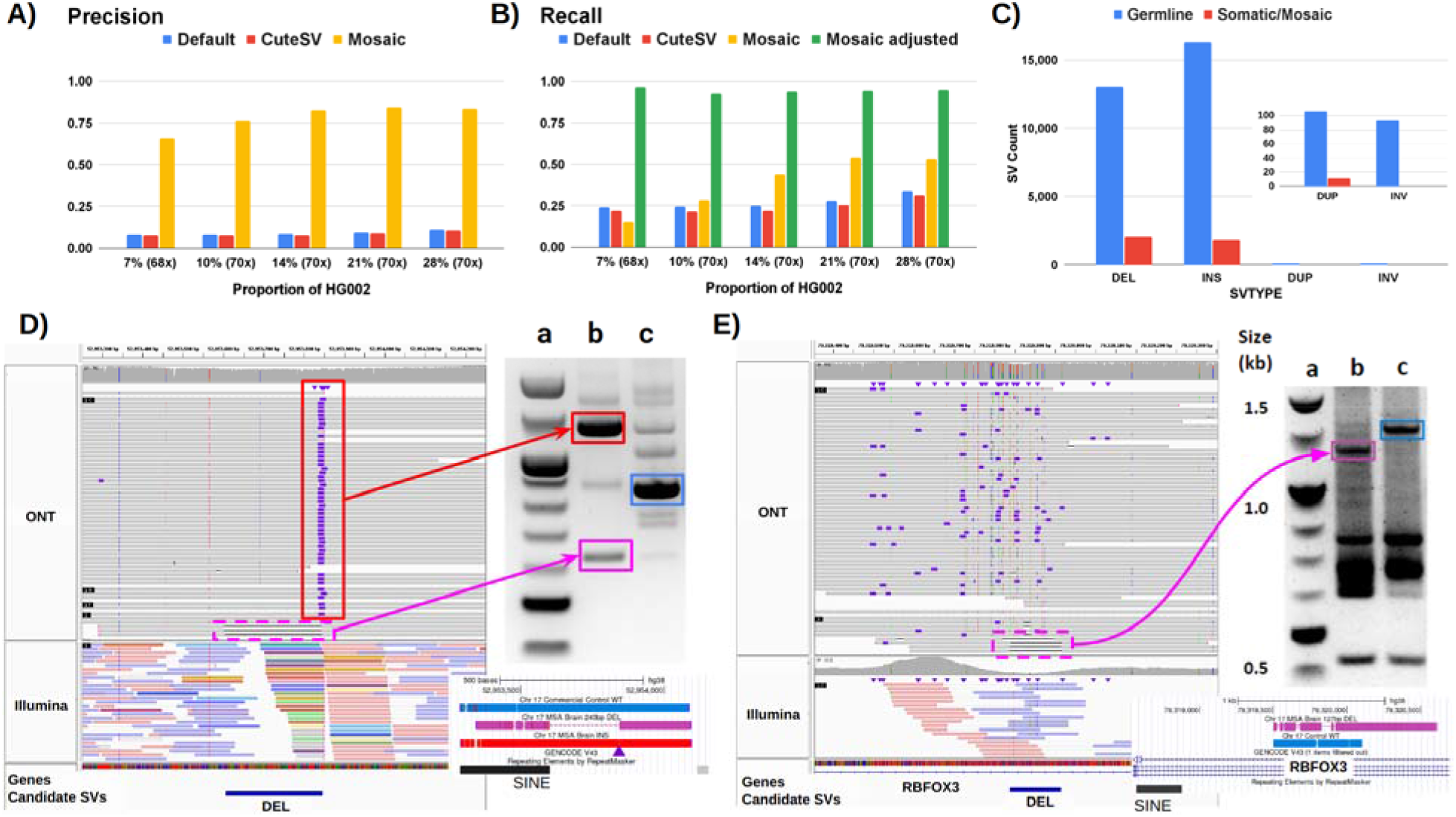
Recovery of somatic SVs using the Sniffles2 non-germline mode. **A+B)** benchmark of mixtures of HG002 with HG00733. We spiked HG002 in various concentrations and measured the precision (A) and recall (B) of Sniffles2 default (blue) and mosaic modes (red), alongside cuteSV (in yellow). For the recall we added an adjusted recall (in green) as Sniffles2 mosaic mode only call SV in the range of 0.05 to 0.20 VAF, and thus everything outside that range will not be analyzed. **C)** Overview of the number of SV types identified as germline (blue) and mosaic in the cingulate cortex brain region of an MSA patient brain sample sequenced with 55x ONT long-reads. A zoom is shown for Duplication and Inversion SVs. **D+E)** Validated mosaic SVs detected by Sniffles2. **D)** mosaic deletion close to a germline Alu insertion. The IGV screenshot shows bulk whole genome sequencing: top panel 55x ONT bottom panel 85x Illumina. PCR validation shows both products from the MSA brain (column b, insertion in top and deletion in bottom) compared to a control (c) and the ladder (a). The PCR products highlighted in squares were Sanger sequenced, and the alignment is shown below the gel (colors matching), with the INS position marked with a purple triangle. **E)** mosaic deletion within RBFOX3. The IGV screenshot shows bulk WGS: top panel 55x ONT, bottom panel 85x Illumina. PCR demonstrates the mosaic deletion (column b, WT in top and deletion in bottom) compared to two controls (c, brain control) and the ladder (a). The PCR products highlighted in squares were Sanger sequenced and the alignment is shown below the gel (colors matching). **Supplementary figure 9** shows the complete unannotated gels, **Supplementary figure 10** shows a different view of the same Illumina results for panel **4E. Supplementary** table 14 shows the complete list of candidate SVs and **Supplementary figure 11** shows all IGV screenshots for the same candidates.

Next, we applied Sniffles2 mosaic mode to an affected brain region (cingulate cortex) of an MSA patient at 55x coverage using ONT. Here we are interested in all types of SVs, including rearrangements (INV & DUP). In this particular case, however, we need to be alert for the possibility that chimeras can form inversions or other duplications and as such contribute to the overall apparent somatic SV calls. To avoid this Sniffles2 deploys filters for low frequency inversions that are 1kb or smaller. Figure 4C shows the overall number of SV and their type for both the germline and mosaic SV call-sets (**Supplementary table 13**). We detected a higher proportion of deletions than insertions in the mosaic calling when compared to germline (INS/DEL ratio 1.3669072 germline, 0.7760922 mosaic). We compared this ratio across all samples used in the paper and found an average INS/DEL ratio of 1.10206 for germline SV calling. Thus clearly showing differences between mosaic and germline SV. **Supplementary table 14** shows 34 mosaic SVs that were manually curated and Figure 4D and **4E** show two of them, both deletions, which were validated by PCR and Sanger sequencing. One is overlapping a repeat element, and one affects a neuronal gene. Figure 4D shows an example of a mosaic deletion close to a germline insertion that was identified using 55x ONT long-reads (top IGV panel). Interestingly we observed these events were located between Alu elements, one novel insertion and one already pre-existing on the reference. We compared the insertion sequence to the neighboring Alu sequence and found great similarity (89.17%). This particular case is a direct orientation of an AluY, which is the Alu subfamily that is most predisposed to brain recombination and thus leading to mosaic deletions^61^. We then performed a blast search of the insertion sequence reported by Sniffles2 (and Sanger sequencing, 100% identity) and found that it belongs to an AluYa5. Across this sample we could identify 25 other regions that had similar alleles of Alu insertions that lead to mosaic deletions both identified with Sniffles. When expanding our search to other sizes of insertions we identified a total of 206 regions where insertions might lead to an instability of the region causing a mosaic deletion in the proximity.

This again highlights the ability of Sniffles to recover potential interesting alleles genome wide and at scale. Figure 4D further shows discordant Illumina reads (colored) indicating multiple translocations instead of the actual Alu insertions, which we reported previously^29^.

**Figure 4E** shows another example of a mosaic deletion, this time overlapping an intron of the *RBFOX3* gene which encodes NeuN, an important nuclear antigen used for sorting neuronal nuclei^62^. On manual inspection of the short reads (**Supplementary Figure 10**) we observed this deletion also on the Illumina reads (5 reads out of ∼85x), but was not identifiable using Manta. Figure 4D and **4E** also show the result of PCR validation of both SVs. For the first validated SV (**4D**), the PCR gel shows both the insertion and deletion event (column b) with the proper SV length of 240 bp reported by Sniffles. For the second validated SV (**4E**), the PCR gel shows evidence of the 127 bp deletion. We further validated these SVs by Sanger sequencing the PCR products highlighted in both gels, which again showed both deletions detected by Sniffles.

Next, we assessed the potential of the identified mosaic SV from Sniffles impacting genes in this sample. We identified 3,049 Sniffles2 non-germline SVs that have read support unique to the cingulate cortex region from which. 2,856 of these SV overlapped with 1,176 genes including at least 80 that are related to the brain and its development (**Supplementary Table 15**). Some examples are *GABRB3* which is implicated in many human neurodevelopmental disorders and syndromes, *NRXN3* which is involved in synaptic plasticity and Netrin receptor DCC which is involved in neuron migration and axonogenesis. *GRIN2A*, which encodes for *NMDA* receptor subunit 2a. Of these 29 SVs, 27 overlapped introns and 2 affected at least one exon. Furthermore, 4 SVs disturb regulatory regions associated with at least one gene, including *PEX26*, *DLL1* and *ABCA2*. This further highlights Sniffles2’s ability to not only detect SVs that cannot be identified using Illumina data alone but also the likely unique presence of a subset of these calls within a brain region. Their distribution across functional areas of the genome in a brain affected by multiple system atrophy merits further study.

A significant fraction of germline SVs is known to be associated with genomic repeats such as *Alu* and L1 elements. To understand the differences between germline and non-germline SVs in this regard, we separately analyzed the association of Sniffles2 germline and non-germline SV calls with different repeat families (**Supplementary Table 16**, Figure 5C). In summary, we found a similar fraction of germline and non-germline SVs was associated with repeat elements (60.61% and 59.70% respectively). When comparing specific repeat elements, we found Alu being more abundant in germline SV (4.82% difference) and simple repeats more abundant in non-germline/mosaic SVs (6.54% difference). Interestingly, the patterns of repeat association only shifted in duplication where LTR, L2 and other repeats differed from the norm and with inversions where no SV was detected in the mosaic SVs (Figure 5C and **Supplementary Figure 8**). For duplications, the fraction of non-germline SVs associated with LINE1 and simple repeats showed the highest difference over germline SVs. This highlights the different ways in which repeat elements are associated with somatic structural variants. As we could show above Alu insertion mediated mosaic deletions. **Supplementary Figure 12** shows two examples of read alignments for a non-germline duplication SV that was solely called by Sniffles2, alongside with its relation to nearby and overlapping repeat elements. For deletions and insertions, we observe similar to slightly lower fractions of non-germline SVs associated with most repeat types, with the exception of simple repeats. For this repeat family, especially non-germline duplication and insertion SVs had a higher fraction associated. Repetitive elements may be associated with neurodegenerative disorders, through increased expression and / or de novo somatic genomic integration^41^. The observance of a higher fraction of non-germline insertion and deletion SVs being associated with simple repeat elements could suggest a further correlation for this on the level of an individual brain region. Overall, this also highlights the differences between repeat families in their effects on somatic SV generation^63^.

**Figure 5:**
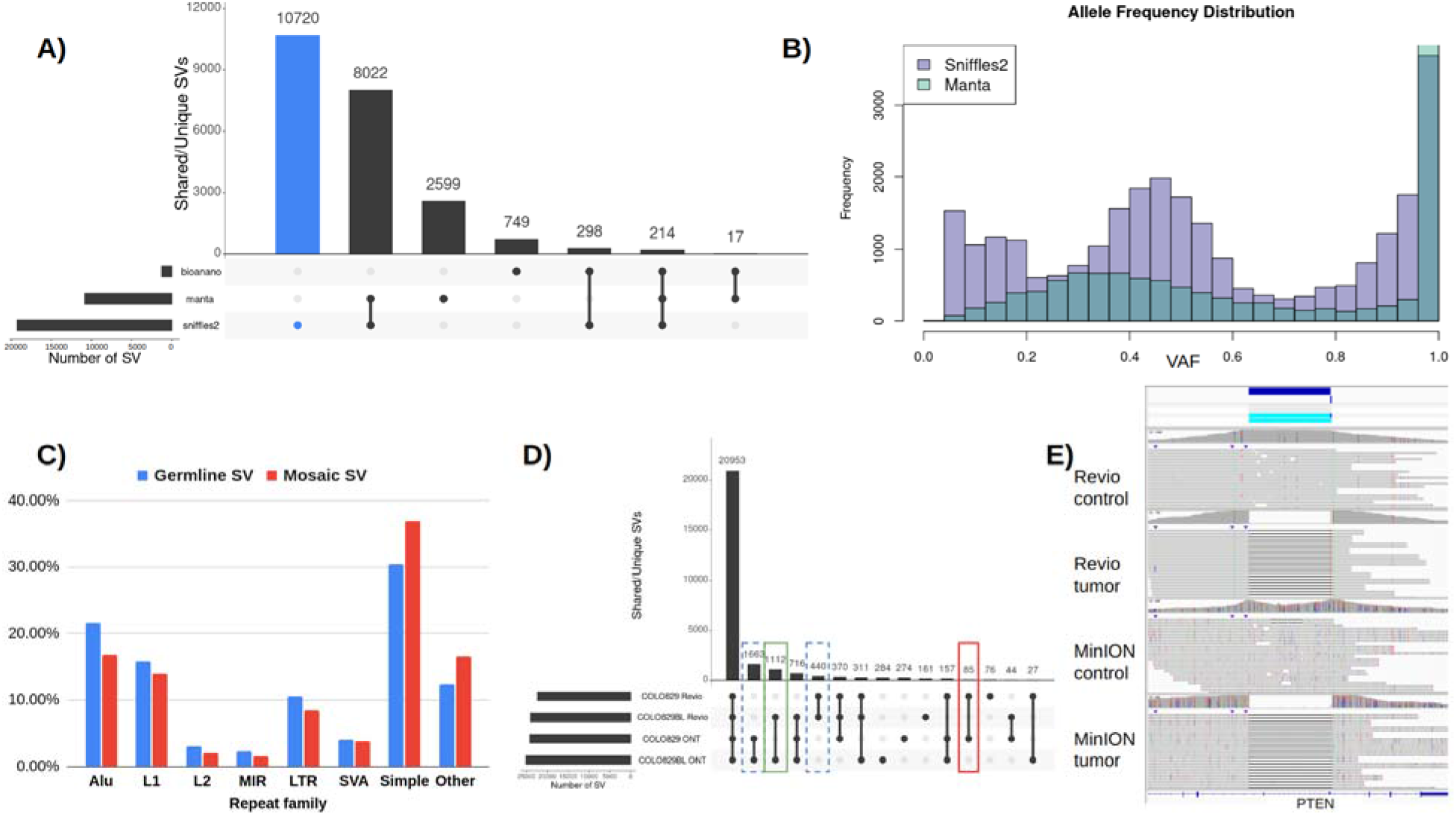
Insights into somatic SVs in the MSA patient brain sample. **A)** Overall comparison of SVs detected in ONT (Sniffles2), Illumina (Manta) and OGM data sets. **B)** Distribution of allele frequencies for SVs identified by Sniffles2 and Manta. **C)** Association of Sniffles2 germline and non-germline SVs with repeat elements. **D)** Tumor-normal comparison of the COLO829 cell line using two different sequencing technologies, ONT MinION and Pacbio Revio. Highlighted are the tumor specific SVs (in red), the normal/control specific SVs (in green) and the technology specific SVs (dashed lines). In the cancer specific SV we can find variants overlapping with cancer related genes such as PTEN, PMS2, ARHGEF5, PAK2, WWOX. Differences between ONT and Revio calls for the same cell-line can be attributed to either technology differences and the evolution of the cell-line through time. **E)** Example of a cancer-specific somatic SV that affects the PTEN gene. Both the Pacbio and ONT dataset showed the same coordinates for the variant and no read support is found in the control.

Next, we compared the different technologies to the Sniffles2 results. The same brain region was also sequenced by Illumina short reads (90x) and analyzed by Bionano optical genome mapping (OGM 690x) (see methods). The Sniffles2 calls from both germline (21,965) and mosaic (2,937) were concatenated as they VAF are mutually exclusive. For illumina, Manta^64^ detected 12,142 SV, and OGM (5kb or larger) 1,463 SVs. Figure 5A highlights the agreement of all SVs detected in the same sample by the three technologies for a minimum length of 50bp and excluding translocations (**Supplementary Table 17.1**). From the total of 22,619 merged SV calls, only 214 are shared across all methods and 19,254 (85.12%%) were detected by Sniffles2 together with either Manta, OGM or both. Sniffles2 uniquely detected 10,720 (47.39%) SVs, while Manta uniquely detected 2,599 (11.49%) and OGM 749 (3.31%). We next identified the most abundant SV type per method for the unique calls. We found that Sniffles2 has a higher number of insertions and deletions (94.87%, 70.98% respectively%), while for Manta are duplications and inversion (98.35%, 92.12% respectively, **Supplementary table 17.2**,). At the same time, Illumina showed a higher overlap with Sniffles2 (8,022 SVs) compared to OGM (298 SVs), likely also because of the minimum length for OGM.

The overlap between Sniffles2 and the other technologies was also much higher for deletions (4,801) than for insertions (3,607). These differences are likely explained by the individual difficulty in detecting larger (i.e., Illumina) and smaller (i.e., OGM, recommended threshold approximately 5kbp or larger) insertion events, respectively. Next, we took a closer look at putatively mosaic SVs detected by Sniffles2. For this, we separated Sniffles2 detected SVs by their reported variant allele frequencies (VAF) into germline (VAF > 0.3) and likely mosaic (VAF ≤ 0.2) calls. Interestingly, a total of 58 insertions were shared uniquely by the germline and mosaic calls of Sniffles2. No other SV types were observed overlapping between mosaic and germline. Interestingly we do observe size differences for these 58 SV but share the same start location.. Next, we compared the mosaic Illumina and OGM calls. Only 702 mosaic SVs reported by Sniffles2 could be detected by either Illumina or OGM or both (24 OGM, 666 illumina and 12 both), highlighting the difficulty in identifying rare SVs. For deletions, only 473 of SVs were also found by either OGMand/or Illumina. Only 218 mosaic insertions reported by Sniffles2 were detected by the other methods. For duplications and inversions, only the Illumina data showed overlap (one and ten, respectively) with Sniffles2 mosaic SVs.

We further noted a shift in the allele frequencies across the Manta calls compared to the Sniffles2 calls (Figure 5B). As expected, for Sniffles2, we observe a multimodal distribution with three peaks, representing homozygous, heterozygous and non-germline SVs, respectively. In contrast, Manta shows two main peaks in their allele frequency distribution. A homozygous (∼0.9-1 AF) and a broad peak around 0.3 AF, which would be below the typical expected heterozygous AF peak. For Sniffles2, we furthermore observe a ∼140% increase in the area under the curve in the putative non-germline range of allele frequencies (0.0-0.2) when compared to the illumina data, thus showing the potential of Sniffles2 to detect low frequency SV.

Finally, we focused on the non-germline (mosaic/somatic) SVs exclusive to the cingulate cortex brain region. For this, we also sequenced the neighboring cingulate white matter from the same patient using Illumina. We used SVTyper^65^ to genotype Sniffles2 SVs (only deletions, duplications and inversions) that were not initially identified by Manta against the aligned Illumina reads from both brain regions (**Supplementary table 17.3**). This way we identified 497 SVs that initially were not identified in Illumina, but were genotyped as present. We identified 484 non-germline SVs using Sniffles2 that have Illumina read support in the neighboring brain region. Thus, showing that Sniffles2 is able to accurately detect low-frequency (ie mosaic) SVs.

### Identification of cancer-specific somatic SVs by Sniffles2

We have shown that Sniffles2 can accurately identify SVs across the whole range of allele frequencies. Furthermore, we showed that Sniffles2 enables fast and accurate population SV calling. Next, we made use of the population merge strategy of Sniffles2 to investigate its ability to identify tumor-specific somatic SVs using paired tumor/normal samples. For this endeavor we used the well-known and highly studied COLO829 cancer cell line (with blood control COLO829BL). By merging the SV calls from both tumor and normal we are able to identify tumor-specific somatic SVs based on the genotype, alternative read count and and the support vector. Figure 5D shows shared and unique SV between the COLO829/COLO829BL cell-lines for two long-read technologies: ONT MinION and Pacbio Revio.

First, we tested both germline and mosaic SV calls against a COLO829 benchmark dataset^66^. This benchmark was done by analyzing the COLO928 sample using multiple technologies. We filtered out all SV smaller than 50 bases, and divided the analysis in two parts: translocations (13 BNDs in the benchmark) and the rest of the SVs (49 SVs in the benchmark). For the case of the translocations (BND) Sniffles2 reported five BND, from which four are present in the benchmark with all within five bases of either breakpoint. Nine BNDs from the benchmark were not detected by Sniffles2, and one BND was called but not present in the benchmark (**Supplementary Table 18**). For the rest of the 49 SVs (INS, DEL, INV, DUP), Sniffles2 was able to identify 36 in either the ONT, Revio datasets or both with breakpoints within 300 bases (max. allowed distance between breakpoints = 1kb, **Supplementary Figure 13**), moreover in some cases the technologies didn’t agree on the genotype (het vs. homozygous alt.). This highlights potential changes across the cell lines as the ONT and Pacbio data were released a few years apart. From the remaining unidentified SV, four had no reads for the alternative allele (0% AF), three were identified by Sniffles as mosaic (8.2-21% AF), four had heterozygous genotype (31-41% AF) and two had homozygous alternative genotype (81-100% AF, **Supplementary table 18, Supplementary figure 14).** For Sniffles2 it shows that we missed only six germline and three mosaic SVs from the benchmark (81.63% recall, **Supplementary table 18**). These results show the difficulty of defining a benchmark dataset for an ever-evolving cancer cell-line such as COLO829 or any other.

We further annotated 79 tumor specific somatic SVs identified by Sniffles (see Methods). These SVs overlap with cancer related genes such as *PTEN, PMS2, ARHGEF5, PAK2, WWOX* to mention some. Additionally, some SVs overlap with olfactory receptor pseudogenes. Figure 5E and **Supplementary Figure 15** shows examples of cancer-specific germline SV. Manual inspection in IGV identified homozygous, heterozygous and LoH events.

## Discussion

In this paper, we present a new version of the highly popular SV caller: Sniffles. Sniffles2 is a significant improvement in terms of accuracy and runtime not only compared to Sniffles v1, but also to all other commonly used long-read based SV callers (see Figure 2). We show higher accuracy across different coverages (5-50x) using different sequencing technologies (Pacbio HiFi and ONT) and even across all SV types. This is achieved by an automatic parameter optimization that is part of Sniffles2 compared to all other SV callers that require manual adjustments. Besides this Sniffles2 is also able to genotype SV and leverage phased reads (using HP and PS tags) as input to provide phased SV in a VCF file. Sniffles2 is more than a simple extension (Figure 1). For the first time, we demonstrate a gVCF concept for SV calling and implemented a working version in Sniffles2. This instantaneously halves the requirements of computing and storage for population/family SV or even tumor vs. normal SV calling. Thus resolving the ever larger demands of long-read data sets^21^. Furthermore it solves the n+1 problems when a new sample is added on a later stage of the project. We demonstrated the new merge concept across the GIAB family where Sniffles2 produced a fully genotyped VCF file within minutes resulting in a low Mendelian error rate and negligible missing genotypes. We could demonstrate the utility across the 31 ONT Mendelian samples, where Sniffles2 resolved SVs mapping to complex regions of the genome with a direct impact to disease, supporting copy-number data. This clearly illustrates the benefit of this novel approach that can easily scale to new population long-read challenges. Moreover, this strategy can also be applied to sample comparison like in the example shown of tumor-normal analysis to detect cancer-specific somatic SV. Currently, there are in development other somatic SV callers^67^ that are specialized on tumor vs. normal tissue comparison. In contrast to them Sniffles2 is a general purpose SV caller that can also be used to detect cancer-specific somatic, but further mosaic SV. Furthemore, the same strategy can also be used to compare different tissues within the same organism (for example different brain regions)

Another novelty of Sniffles2 is the mosaic-mode that enables the detection of low- frequency SVs with a standard sequencing run, while maintaining a high precision. We have demonstrated this novel approach using a synthetic data set of HG002 and a genetically unrelated individual HG00733. This showed the accuracy and recall of Sniffles2 while depending on only 2-3 reads overall to distinguish SV from noise (Figure 4 **A and B**). We then turned our attention to MSA, a rare sporadic neurodegenerative disease related to Parkinson’s, with negligible heritability (<7%)^68^. We performed ONT WGS on an affected brain region from one patient, where Sniffles2 was able to identify presumptively low-frequency mosaic SV and showcased great performance partially validated by Illumina and optical genome mapping approaches, thus overall highlighting the fact that Sniffles2 is highly versatile and accurate. While thresholding on the variant allele frequencies (here 5-20% AF) for the identification of potential somatic variants is straightforward, there is still a gray area to be addressed. For multiple SV, we saw a continuum in VAF (between 20 and 30% AF) which suggests that some SV with apparent AF < 30% may also be germline. Thus, the comparison to population data or to different tissues is still favorable (e.g. tumor vs. normal). It is interesting to note that the detection of tissue specific SV as proposed here can be impacted by multiple biases. First we can have a detection bias in the Illumina data (e.g. insertions) but further a sampling bias in the other tissue might also result in tissue specific SV detection. The possible role of somatic SV’s in MSA is under investigation^44^, although further validation data from more cases and controls would be required to allow interpretation of the present findings. Interestingly, in our experiments at mosaic level, we identified many more deletions than insertions in contrast to germline (AF>0.2). We speculate that this is indeed a biological signal and not a detection bias due to NAHR or other mediated mechanisms.

Despite all the novelties and solving central problems of SV calling at scale and accuracy for long-reads many challenges remain. We still lack high quality benchmarks for non-insertion and deletion calls including complex rearrangements. Achieving this will boost the field of SV detection further and promote novel methodologies. In this study, we could only evaluate other SV types (inversions, duplications, and translocations) via simulated data using an established pipeline. **Supplementary Figure 3** summarizes these results. While this is unsatisfactory, it remains the only way as benchmark sets focus on insertion and deletions only. In addition, Sniffles2 does not yet solve the issues with highly rearranged regions where SVs can be overlapping to each other. This remains a near future goal of Sniffles2 and will also require improved benchmark sets and even standards to report these events as the VCF standard does not provide a clear recommendation. Currently these complex alleles would need to be reported as indepentend BND events, which looses their individual impact (e.g. DUP/ INV-DUP) on the region itself. Nevertheless this is clearly needed as our experiments on the Mendelian cohort shows. Another highly important factor that Sniffles2 improves is tumor normal comparisons. Here, we included an example of using the population merge to detect cancer-specific somatic SV, moreover we are still working on how to best address the detection of the low variant frequency SV which are common in cancer.

Overall this paper reports the innovations across Sniffles2 and highlights them across Mendelian cases, an MSA patient and a tumor-normal comparison. We believe that our new implementations (merging and mosaic mode) will spark novel findings across human diseases and diversity. Furthermore we believe that these will also be important for other species. Despite the fact that the genotype model for Sniffles2 is designed for diploid organisms, Sniffles2 is capable of also detecting SV in haploid (as shown for X chromosomes in males) or polyploid organisms. For higher ploidy levels we would suggest running the mosaic mode as otherwise the genotype caller will penalize true SV. Thus, again highlighting Sniffles2 as a highly accurate and versatile method to detect SV of any kind and property.

## Online Methods

### Patient Enrollment

The 31 individuals (proband and parents) included in this study were enrolled into research protocols approved by the Institutional Review Board (IRB) at Baylor College of Medicine and the Pacific Northwest Research Institute (H-29697 and H-47127, WIRB#20202158).

### Sniffles2 methodology

#### Sniffles2: Germline calling

An overview of the steps involved in Sniffles2 germline SV detection algorithm is shown in **Supplementary Figure 16.**

Sniffles2 germline mode accepts aligned long-reads as input (BAM or CRAM format, sorted by genomic coordinate and indexed). First, read alignments are parsed and pre-filtered based on minimum mapping quality (default: 20), minimum alignment length (default: 1kb), and maximum number of split alignments (default: 3 + 0.1 *ReadLengthKb*). Split alignments are analyzed to extract SV signals for insertions, deletions, duplications, inversions, and breakends. Next to analyze splits, inline alignments are scanned for insertion and deletion signals. Sniffles2 does not merge nearby inline insertion and deletion events at this point. SV signals that fulfill a minimum length threshold (default: 0.9 *MinSVLength*) are subsequently recorded in high-resolution genomic bins. Start and end positions of alignments are recorded in a separate data structure for facilitating later coverage computation without requiring reopening of alignment files.

Sniffles2 employs a three-phase clustering process to translate individual SV signals into putative SV candidates. First, SV signals extracted from reads in the preprocessing step are clustered based on their indicated SV type and genomic start position. Second, insertion and deletion sequences in each cluster stemming from the same read are merged to correct for alignment errors in highly repetitive regions. Third, preliminary clusters are re-split to represent different supported SV lengths.

The first clustering phase constitutes a fast pass over all bins (default bin size: 100bp) containing SV signals extracted from alignments in the preprocessing step. Bins are traversed from chromosome start to end separately for each of the five basic SV types. Neighboring bins are merged if the inner distance between them is smaller than a threshold calculated based on the minimum standard deviation of the genomic SV start positions within each bin. The inner distance threshold d_n_ is calculated as: *d_n_* = *r* . *min*(*σ_StartA_*, *σ_StartB_*), where r is a constant (default: 2.5), and *σ_StartA_*, *σ_StartB_* refer to the standard deviation of indicated SV start positions in the two neighboring bins, respectively. In regions spanning tandem repeats, a more relaxed clustering criterion is applied: Neighboring bins are also clustered when their outer distance falls below a threshold defined based on the indicated average SV length of the SV signals stored in the neighboring bins. This threshold d_r_ is calculated as: *d_r_* = *min*(*h_ffiaS_* , *h* .[*x_A_* + *x_B_*]) , where h and h_max_ are constants (default: 1.5 and 1kb, respectively) and x_A_,x_B_ refer to the mean indicated SV length in the two neighboring bins. Whenever two neighboring bins have been merged, the clustering is restarted at the bin preceding the merged pair, facilitating the growth of SV clusters in both upstream and downstream directions. The first clustering phase is completed as soon as the last bin in the chromosome has been reached.

The second clustering phase constitutes merging of insertion and deletion events stemming from the same read that have been placed within the same initial cluster. Events with an inner distance closer than the set threshold (default: 150bp) are merged. In areas of tandem repeats, the distance threshold is set to the size of the initial cluster itself.

In the third phase, clusters are split by indicated SV length of the contained SV signals and subsequently re-merged, which leads to the final separation of SVs that share a start position on the reference but have different lengths. Bins are traversed from those containing small to large SV signals and merged in a similar fashion to phase one, based on the relative difference in SV length between neighboring bins being no larger than a given threshold (default: 0.33). In clusters overlapping tandem repeats, Sniffles2 does not perform resplitting.

Differentiated clustering parameters are applied to breakend-type SVs, since no length is available as a metric to drive clustering.

At the beginning of postprocessing, SV candidates are generated from the final clusters resulting at the end of the last stage. Start coordinates and SV length are determined based on the median of the most common values supported by the reads. Standard deviations are calculated for the trimmed distribution of indicated SV start position and lengths. The quality value is summarized as the mean mapping quality of supporting reads. SVs are labeled as precise when the sum of SV start and length standard deviation is less than the set threshold (25bp).

SV candidates are filtered based on absolute and relative (compared to the SV length) standard deviation of their coordinates. In addition, type-specific coverage filtering is applied to deletions and duplications, requiring central coverage changes consistent with the detected variant. Instead of requiring users to settle for a predefined, static minimum read support threshold, Sniffles2 dynamically adjusts the minimum support value based on estimates of global and regional sequencing coverage. By default, the minimum read support threshold is calculated as *MinSuppart* = *a* . ([1 - *λ*] *Cglabal* + *λ Clacal*). Where Cglobal and Clocal refer to average chromosomal and SV surrounding coverage, respectively. The parameters are set as *α*=0.1 and *λ*=0.75, by default. For insertion and deletion SVs, support from inline alignments and split alignments is output separately. Additionally indicated support from soft-clipped reads is additionally recorded for insertion SVs.

Genotypes are determined using a maximum-likelihood approach. The genotype quality is calculated based on the likelihood ratio of the second most likely to the output genotype: *Q* = -10 *log*_10_(*L*_2_/*L*_1_), whereas L_1_ and L_2_ refer to the likelihood of the most likely genotype and second most likely genotype, respectively. Genotype likelihoods are computed for a binomial distribution for the observed number of variant and reference reads. Genotype likelihoods are set as 1.0-ß for 1/1, 0.5 for 0/1, and ß for 0/0, whereas ß represents the genotype error introduced through sequencing and alignment artifacts and is set to ß=0.05 by default.

For insertion SVs, sequencing and read aligner errors are corrected using a fast kmer-based pseudo-alignment method. Through this, Sniffles2 generates a consensus sequence in two steps: In the first step, the best possible starting sequence is chosen from the supporting read with the smallest distance in SV start position and length to the final reported SV coordinates. K-mers (default length: 6bp) are enumerated for this reads supported insertion sequence and a taboo set of repetitive k-mers, which occur more than once in the sequence, is built. Simultaneously, the positions of non-repetitive k-mers are stored in an anchor table to facilitate pseudo-alignment of the other reads. In the second phase, k-mers from other reads insertion sequences are enumerated. When a kmer is present in the anchor table, the corresponding position in both the initial insertion sequence and the current read are stored. After all reads have had their k-mers anchored, sequences between anchored kmers are extracted from the pseudo-aligned reads. These sequences from between the anchored kmers constitute the parts of each reads insertion sequence in disagreement with the initial sequence. Finally, coordinates of the initial sequence are traversed and the consensus is generated as the most common base at the respective position throughout all pseudo-aligned reads. Long insertions (ie. multiple kbp) are often difficult to detect even in long-read data because reads often do not span the full insertion sequence. To improve detection of long insertions, Sniffles2 records these clipped read events as additional support for presence of a large insertion. This enables Sniffles2 to accurately detect large insertions even when the SV is fully covered just by a single read.

Post processed and annotated SV calls that passed quality control checks are written to the output VCF file. Quality control filters applied to SV candidates by default include absolute and relative standard deviation of the SV breakpoints, coverage change for copy number variants and minimum coverage in the surrounding genomic region. Additionally, all unfiltered SV candidates and genome-wide coverage information are written to a specified output SNF file, which may be consecutively used as input for multi-sample calling (see Section 1b). Using the -- qc-output-all option, all unfiltered candidates (except for the minimum SV length filter) can also be directly written to the VCF output file complete with the respective reasons for why they would have been filtered by default.

Full parallelization across chromosomes is applied through all key steps in Sniffles2, including preprocessing, clustering and postprocessing. The final SV calls are written to a sorted VCF output file. Alternatively, Sniffles2 also supports direct output to a sorted, bgzipped and tabix-indexed VCF file.

#### Sniffles2: Combined Calling (Population Mode)

Sniffles2 produces a fully genotyped population VCF file by introducing a specialized mode (*Sniffles2 combine*) for both family- and population-level SV calling. *Sniffles2 combine* is built around a new specialized binary file format (SNF), designed to store a complete snapshot of structural variation and sequencing coverage for a single sample. Mergeable SNF files for later population-level calling are designed to be easily produced as a side-product of regular single-sample SV calling using Sniffles2, by using the optional --snf output argument. Based on individual use case requirements, Sniffles2 can simultaneously produce SNF files and/or regular VCF files in a single run of processing an individual sample.

SNF files consist of a JSON-based index followed by a series of multiple gzip-compressed blocks (separated by genomic coordinates). Each block stores all putative SV candidates, separated by SV type, for a single sample’s respective genomic region. This includes candidates only supported by e.g. a single read, that would normally be ignored. Each block furthermore stores sequencing coverage information (500bp resolution by default). All stored SV candidates contain a compressed form of all the information of the final SV calls, as they would be output in a single-sample VCF file, such as start, end positions, standard deviation and alternative alleles. SNF blocks span a genomic region of 100kb by default. This small block size comparison to a typical mammal genome allows Sniffles2 to combine a high number of samples simultaneously while keeping a manageable memory footprint.

SNF files, once generated, can then be used as input for the *Sniffles2 combine* mode, producing a final, fully genotyped population-level VCF file within seconds. SNF files may also be reused in the combine step, e.g., when the population is later on extended, when individual samples need to be re-run, or when querying whether a later newly identified SV is present in a population. These use cases would not be possible without costly reprocessing of all samples with the currently prevalent method of forced calling. A schematic of SNF file structure can be found in **Supplementary Figure 17.**

When presented with multiple SNF files as input, Sniffles2 combines them through a single pass over chromosomal regions. For each region, the respective SNF blocks overlapping it are loaded, including all SV candidates and coverage information from each sample. In the following step, Sniffles2 groups the loaded SV candidates based on SV type and coordinate-based matching criteria. For each SV candidate, Sniffles2 first checks if there is an already existing, matching group. An SV candidate matches a group if it has the same SV type and the sum of start position and length deviation is less than 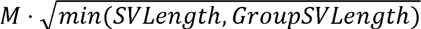, where M is set to 500bp by default (user-adjustable). The start position and SV length of a group are defined as the arithmetic mean of all SVs currently contained in it. In case there are one or more groups that fulfill the matching criteria for the current SV candidate, the group with the smallest deviation metric is chosen and the SV candidate placed therein. The coordinates of the selected group are then subsequently updated to represent the new average position of length of the contained candidates. If there are no matches, a new SV group is created. By default, Sniffles2 allows for matching multiple SVs from the same sample within a group (can be disabled using *a* dedicated parameter).

This partition of SNF files into individually loadable blocks keeps Sniffles2 memory footprint manageable even when processing a high number of samples and/or samples with high-coverage. Sniffles2 further implements a dynamic binning strategy for accelerating the grouping phase. Sniffles2 first assigns all loaded SV candidates from the current chromosomal region to bins based on SV type and start position. Bins are then traversed from low to high coordinate within the current block, while collecting encountered SV candidates. When the number of SV candidates exceeds a certain threshold (default: *PapulatianSize* * 0.5), the collected SV candidates are grouped as described above. Triggering the grouping stage only when a set number of SV candidates is reached allows for the highest possible accuracy in matching SVs from different samples in regions with low complexity, while keeping the runtime manageable even in regions with a high density of SV candidates. To avoid edge effects, the final resolving of SV groups with genomic coordinates close to the ends (default: <2.5kb) of the respective bin are carried over and finally resolved in conjunction with the grouping of the next bins. The same strategy is applied to SV groups close to the genomic start or end coordinate of the currently processed SNF block.

By default, Sniffles2 combine mode will output all resulting SV groups in the population that meet at least one of two criteria:

A. The SV has been detected with high-confidence (i.e. passes all quality control checks) in at least one sample and/or
B. By default, to have a high-confidence call in at least one sample.

The SV is present in a sufficiently high number of individual samples, even though it may not have passed individual quality-control checks (default: present in at least *max*(0.2 *PopulatianSize*, 2) samples). These parameters are also user-adjustable and can be adjusted or disabled without having to re-generate the SNF files for the individual samples.

Each final SV group that passes the above criteria is output as an SV in the final population-level VCF file, including the genotypes from all samples. For samples that did not have a SV candidate that could be matched to the group, Sniffles2 first uses the coverage information stored in the SNF file of the respective sample to determine if the sequencing depth at around the group’s genomic location was sufficiently high (default value: 5x). If it is, the sample genotype for that SV is output as 0/0 if there is no evidence, and otherwise as missing (./.). For all SVs, the number of reads supporting the SV and supporting the reference are output for all samples, allowing for differentiation between true biological and technically induced absence of each SV from a sample.

Sniffles2 combine is fully parallelized, allowing leveraging multi-core CPU systems not just for calling individual samples but also the final combination step. This, in conjunction with the separation of SNF files into blocks and dynamic binning strategy, together enables Sniffles2 to perform scalable population-level SV calling.

#### Sniffles2: Low frequency SV (mosaic) calling

In the non-germline mode, a reduced default minimum support multiplier is applied (default: 0.025) to increase sensitivity for low-frequency SVs. At coverage levels of 30x to 50x, this leads to a minimum read support of 2-4 reads for the detection of non-germline SVs. To balance out the increased influence of sequencing and alignment artifacts at this lowered read support threshold, additional filtering based on alignment quality is applied. In the preprocessing steps, the length-weighted number of mismatches is recorded for all SV signals, excluding insertions and deletions. After calling, SVs with an average weighted mismatch ratio of larger than a threshold *t*= *c* * *a* , where a is the average length-weighted mismatch number for all reads and c is a constant (default: 1.66) are filtered. The additional, coverage-based filtering steps for CNVs applied in the germline mode are not applied in non-germline mode, as coverage changes induced by somatic SVs are not reliably measurable.

### Benchmarking Methodology

#### Computer specifications

All tests are performed in a high performance cluster with Intel(R) Xeon(R) Gold 6148 CPU @ 2.40GHz, the memory allocation is 32Gb unless it is otherwise stated and the number of CPU cores allocated is 8 unless otherwise stated. All CPU time is given as the sum of all compute time as if a single core was used.

#### Benchmarking SV callers on GIAB, 1000 Genomes and CMRG

Reads were mapped using minimap2^69^ (v2.17-r941) technology specific preset parameters. Reference genome GRCh37 was used to test for the GIAB v0.6 SV benchmark and GRCh38 was used to test the Challenging Medical Relevant Genes (CMRG) SV panel. In both cases the ALT and/or Decoy contigs were not included. The -Y option was supplied to disable hard clipping (required by pbsv) and generate the --MD tag (required by Sniffles1) and the PacBio / ONT presets were used respectively. Resulting alignments were converted to BAM format, sorted and indexed using samtools (v1.13).

As measure of coverage across all benchmarked data sets, we used the mapping coverage as reported by mosdepth^70^ (version 0.3.2), which was averaged across all autosomes.

Besides HG002 we also benchmarked SV on three assemblies of the 1000genomes (HG01243, HG02055, HG02080). Here we leveraged the phased HiFi assemblies provided at https://github.com/human-pangenomics/hpgp-data and the corresponding long-reads. The benchmark set was derived from a dipcall^53^ (version 0.2) alignment against the GRCh38 reference. This result was used together with the corresponding bed files for benchmarking.

We used Truvari^51^ (version 2.1) for benchmarking the accuracy of all SV callers across datasets. For benchmarking, we used the *--passonly* parameter to include only those SVs from caller and gold standard that are not marked as filtered. For the GIAB benchmarks, we additionally used the *--giabreport* parameter to generate the benchmark-specific detailed report. As included regions Tier 1 regions were used unless otherwise specified. For all other parameters, default values were used.

Callers were first benchmarked using default parameters and callers other than Sniffles2 were separately benchmarked on GIAB by manually setting the minimum read support parameter to 2 (sensitive).

SVIM^50^ (v1.4.2) does not include filtering steps in its main pipeline, which caused it to perform poorly (F-measure) in most benchmarks, and we were not able to identify a recommended default cutoff for the quality value that SVIM outputs along with its SV calls. Therefore, in line with previous SV caller benchmarks, we filtered the output of SVIM to include only calls with a minimum read support of 10 by default (equal to the default of cuteSV and Sniffles1) or 2 (sensitive).

For benchmarking Sniffles2 (build 2.2), we only used the default parameters with the exception of non-germline SVs, where the *--mosaic* option was supplied. For Sniffles^29^ (v1.12), default parameters were used. For cuteSV^48^ (v1.0.11), we used the additional parameters recommended by the authors for use with HiFi / ONT datasets in their GitHub documentation, as well as the --genotype option. For pbsv^49^ (v2.6.2), we supplied the --ccs option for analyzing HiFi data, as recommended by the authors. Both pbsv and Sniffles2 support the use of tandem repeat annotations for improving SV calling in repetitive regions. For pbsv and Sniffles2, we therefore supplied the tandem repeat annotations for GRCh37 / GRCh38 which we obtained from the pbsv repository on GitHub: https://github.com/PacificBiosciences/pbsv.

For all SV callers that have an option for specifying the number of multiprocessing threads, we set the number of threads as 8. We measure and report the total CPU time and wall clock time using the UNIX time command. For the benchmarks including only insertions and deletions, we used SnpSift ^71^(v4.3t) to filter the output of all SV callers to include only those types of structural variants. To prepare SV caller output for benchmarking, VCF files were sorted using bedtools, compressed and indexed using bgzip and tabix. For SVIM, SVs labeled as INS:NOVEL were relabeled to INS, in order to be able to be matched to insertions in the benchmark sets by Truvari.

#### Simulation of different SV types using SURVIVOR

SURVIVOR^59^(v1.0.7) was used to simulate SV types not covered by the GIAB and other benchmarks. For this benchmark, 3000 duplications, inversions and translocations were each simulated within a length range of 500bp to 30kb on the human reference genome GRCh37 in diploid mode. A total sequencing depth of 30x was simulated for ONT reads, with the error profile obtained using the SURVIVOR scanreads command from the HG002 ONT Q20+ data set. SVs were called using each SV caller for the simulated reads using the default parameters and postprocessing steps also used in the GIAB and other benchmarks (see respective methods subsection). The SURVIVOR eval command was used (matching threshold: 500bp) to obtain TP, FN and FP counts for each caller and simulated SV type from which precision, recall and F-measure were calculated.

#### Measurement of Insertion Sequence accuracy

Accuracy of insertion sequences recovered by the SV callers was measured using Biopython’s^72^ (v1.79) pairwise2 global alignment function. First, the true positive calls from all investigated SV callers on the data set were intersected, to establish a common set of calls to benchmark. Next, the gold standard and reported insertion nucleotide sequences were aligned and the resulting score was normalized by length of the gold standard sequence to compute the alignment identity. We measured sequence accuracy separately for the GIAB HiFi and ONT data sets (30x coverage). Results are shown in **Supplementary Figure 1**. The respective script is made available in the supplementary materials.

#### Simulation of low-frequency SVs

Low-frequency SVs were simulated by combining varying coverage titrations of HG002 and HG00733 into synthetic samples with different levels of mosaicism. Recovery of SVs unique to HG002 was done based on the intersection of SVs of the same type using bedtools with 50% coverage of the SV reciprocally against the truth-set of HG0733^51^. These unique SVs were then used to benchmark to measure recall for low-frequency SVs. For benchmarking the ability of Sniffles2 to detect low-frequency SVs, we simulated synthetic data sets with 63x/5x, 63x/7x, 60x/10x,55x/15x and 50x/20x, where the coverage refers to HG00733 and the second one to HG002. Next, we used the previously selected HG002 unique SVs overlapping the GIAB Tier 1 benchmark. . To measure recall for low-frequency SVs, we ran Sniffles2 in mosaic mode on the synthetic samples and used Truvari as described in the methods section on GIAB benchmarks to compute the recall for the rare HG002 SVs introduced into each HG00733 data set. Simultaneously, we ran sniffles2 and cuteSV with default parameters and benchmarked the results for comparison. Given that sniffles2 mosaic mode only analyzes and reports SV within a defined VAF (5-20%), we excluded all SV that were outside of such VAF to compute an “adjusted recall”. As in all the other GIAB benchmarks, analysis was limited to insertion and deletion SVs.

### MSA patient analysis

#### Optical mapping data on MSA patient brain

Ultra-high molecular weight (UHMW) DNA was isolated from frozen human brain tissues using a Bionano Prep SP Tissue and Tumor DNA Isolation kit (#80038) according to the Bionano prep SP Brain Tissue Isolation Tech Note (#3400). In short, approximately 20mg frozen tissue was homogenized using a Qiagen TissueRuptor (9002755), passed through a 40um filter, and treated sequentially with Qiagen protease (catalog #19155), proteinase K, and RNAse A in lysis and binding buffer. The homogenate was then treated with PMSF to de-activate the Protease and Proteinase K, washed, and eluted. The extracted DNA was mixed using an end-over-end rotator for 1 hour at 5rpm and allowed to rest at room temperature until homogenous (approximately 1 week). 750ng purified UHMW DNA was fluorescently labeled at the recognition site CTTAAG with the enzyme DLE-1 and subsequently counter-stained using a Bionano Prep DLS Labeling Kit (#80005) following manufacturer’s instructions (Bionano Prep Direct Label and Stain (DLS) Protocol #30206). Optical genome mapping was performed using a Saphyr Gen2 platform for a final effective coverage of 894X for the pons and 754X for the cingulate. Effective coverage is defined as the total raw coverage of molecules ≥ 150kbp in length multiplied by the proportion of molecules which align to the reference genome.

Calling of low allele frequency structural variants was performed using the rare variant analysis pipeline (Bionano Solve version 3.6) on molecules ≥ 150kbp in length. De novo assembly was performed using the longest 250X molecules of each dataset. The variant annotation pipeline (Solve 3.7) was used to detect which structural variant calls in the cingulate are present in the pons structural variant calls and/or molecules. See the Bionano Solve Theory of Operations for more details.

#### MSA samples comparison

Illumina reads were mapped to the human genome GRCh38 using bwa^73^ mem (version 0.7.17- r1188) with default parameters including -M to mark split reads as secondary alignments. Subsequently we identified SV using manta^64^ (version 1.6.0).

For ONT, reads were mapped using minimap2 ^69^ (version 2.17-r941) with present parameters for ONT. Subsequently we identified SV using Sniffles2 with both germline (default) and mosaic mode. The Bionano OGM data smap file was converted by SURVIVOR smaptovcf (v1.0.7)into a VCF file.

To compare SVs called by Sniffles2, Manta (Illumina), and OGM (Bionano) we used SURVIVOR merge using a 10kb threshold, matching SV type and ignoring reported SV strand. We extended it to 10kbp after testing 500, 1kbp and 5kp thresholds and observed that the accuracy of the breakpoints from OGM required the larger parameter.

The genotype columns in the SURVIVOR merge output were compared for each SV to determine presence or absence in the results reported by the respective method.

Subsequently, to further investigate SVs absent from the Manta call sets, we additionally genotyped the respective Sniffles2 calls against the raw Illumina read alignments for the same (cingulate cortex) as well as a different brain region (cingulate white matter) using svtyper (version: 0.7.1).^65^ SVs reported as having at least one supporting read by svtyper were considered as present in a sample.

#### PCR validation of selected mosaic deletions

We used NCBI primer design tool to obtain primers straddling the target deletions. The primer sequences for the 240bp deletion were TACCAAGTCTTTCTCCAAGTCCC (forward) and TTGCACAGCCTTGGCTATACTC (reverse) and for the 127bp deletion ATCCTGAGAGAACCCCCTCC (forward) and GGACAGACTCGTGGTTTCGT (reverse). PCR was performed using Phusion Plus PCR Master mix (ThermoFisher), with 0.5µM primers, annealing temperature 60L and extension time 75 sec. PCR results were confirmed using Agilent Tapestation and 2% agarose gel electrophoresis, stained with GelRed (Biotium), with 100 bp DNA ladder (NEB). Initial PCR was performed using 20-40 ng template DNA in 20 μl for 35 cycles. Repeats to obtain adequate products were performed using 100 ng DNA in 50 μl, with 40 cycles for the second deletion, and low-melting point agarose was used to allow relevant amplicon band excision. Extraction and purification from agarose was carried out using QIAquick gel extraction kit (Qiagen). Extracted products, which represented the wild type, deletion, and Alu insertion, underwent Sanger sequencing

### Mendelian inconsistency benchmark in population mode

#### Mendelian benchmark/ inconsistency

To assess the performance of Sniffles2 population mode, we used the Ashkenazim family trio. We called SV using Sniffles2 and cuteSV. For Sniffles2 we used a minimum SV length of 50 and with the output being the SNF binary file that contains the unfiltered SV candidates and genome-wide coverage information (using the --snf option). Then, we merged the SNF files with Sniffles2 population-level calling providing the reference genome to obtain the sequences of the deletions. Here the input are the SNF files and the output the VCF file. For the case of cuteSV we used v1.0.11 with recommended parameters for Oxford Nanopore data (--max_cluster_bias_INS 100 --diff_ratio_merging_INS 0.3 --max_cluster_bias_DEL 100 --diff_ratio_merging_DEL 0.3). Then, we merged the results of cuteSV using SURVIVOR v1.0.7 with a maximum distance between breakpoints of 1kb, a minimum support of one and taking into account the SV type. Next, we performed force-calling with cuteSV using as input the merged SV from SURVIVOR (-Ivcf and --genotype options). Finally, we performed a second merge with SURVIVOR with identical parameters as before.

We then tested the mendelian inconsistency of the genotypes using the BCFtools v1.14 mendelian plugin^58^. The mendelian plugin denotes a genotype consistent when the proband genotype is in concordance with the parental genotypes (e.g F 0/0, M 0/1, P 0/0), inconsistent when the proband and parental genotypes do not match (e.g. F 0/1, M 1/1, P 0/0) and NA when the proband has a missing genotype (./.). For all analysis time was measured utilizing the linux time command.

### Chromosome X disorder patient analysis

Sniffles2 population mode was used to analyze 31 Oxford Nanopore samples that represented cases of Mendelian disorders in probands. We obtained the bam files by running PRINCESS^30^ (version 1.0) using the default parameters and “ont” flag. PRINCESS implicitly calls Minimap2^69^ (version 2.17) with the following parameters “-ax map-ont -Y --MD”; Later, we sorted the output using samtools^58^ (version 1.14). For all samples, unfiltered SV candidates and genome-wide coverage information are written to a specified output SNF file and then merged with Sniffles2 population-level calling. General statistics, such as SV sizes and composition (proportion of each SV type) were computed by extracting the SVLEN, SVTYPE and GT information from the VCF file.

Given the nature of the dataset, only the SV calls from chromosome X were analyzed. Additionally, for specific individuals (BH14379, BH14413) SV from chromosome Y were analyzed given that both aCGH and Sniffles2 called translocations to chromosome Y.

Then, all SVs that were less than 10kb were filtered, as aCGH data showed large events were involved. Finally, we filtered out SV that occurred in the father, as this disorder is fully penetrant in males by comparing the SUPP_VEC tag in the VCF to the sample names. Manual curation was performed for a single SV that was filtered out by the STDEV_LEN filter of Sniffles2 during development.

### Identification of cancer specific somatic SVs by Sniffles2

We used the population level calling (population merge) of Sniffles2 to detect cancer specific somatic SVs by comparing a tumor/normal pair. We used the highly studied COLO829 cancer cell line with the COLO829BL blood control. Structural variants were called with Sniffles2 using default parameters with the --snf option to save candidate SV to the SNF binary file, per sample. We used two tumor-normal pairs, one described in Vale-Inclan et al ^66^, and a sample provided by Pacific biosciences (see Data availability). We then merged the four files using Sniffles2 population merge. Next we analyzed the SV presence/absence by means of the SUPP_VEC tag in the INFO field of the output VCF to extract SV that are only detected in the tumor samples. We compared all the SVs detected by Sniffles2 to the COLO829 SV truth-set to assess the performance of Sniffles2 somatic SV calling. For the case of mosaic SVs we perform the same strategy as before, moreover of the cancer datasets we added the --mosaic option to get the mosaic candidate SVs in the SNF file as well. Here, we also detected somatic SVs but the presence/absence by means of the SUPP_VEC tag in the INFO field to extract cancer-specific SVs.

## Data availability

GIAB HG002 Pacbio HiFi data is hosted at the github server: https://ftp-trace.ncbi.nlm.nih.gov/ReferenceSamples/giab/data/AshkenazimTrio/HG002_NA24385_son/PacBio_CCS_15kb/

ONT HG002: https://labs.epi2me.io/gm24385_q20_2021.10/

ONT HG00733: https://www.internationalgenome.org/data-portal/search?q=HG00733 and https://ftp.hgsc.bcm.edu/Software/Truvari/3.1/sample_vcfs/hg19/li/HG00733.vcf.gz

GIAB benchmark sets:

Genome wide: https://ftp-trace.ncbi.nlm.nih.gov/ReferenceSamples/giab/release/AshkenazimTrio/HG002_NA24385_son/NIST_SV_v0.6/

Medical regions: https://ftp-trace.ncbi.nlm.nih.gov/ReferenceSamples/giab/release/AshkenazimTrio/HG002_NA24385_son/CMRG_v1.00/

The 1000 genomes data sets of the three genomes were downloaded from : https://github.com/human-pangenomics/hpgp-data

The dipcall results that we leveraged as benchmark are deposited at https://github.com/smolkmo/Sniffles2-Supplement

The other data sets have been made available over SRA. 31 Oxford Nanopore data sets that represent cases of Mendelian disorders have SRA bioproject ID PRJNA953021 and dbGaP phs002999.v1.p1. MSA sample has bioproject ID PRJNA985263. The COLO829BL (normal) and COLO829 (tumor) ONT samples can be found with the ENA ID: PRJEB27698 (samples ERR2752451 and ERR2752452 respectively), and the Revio tumor/normal samples can be found in https://downloads.pacbcloud.com/public/revio/2023Q2/COLO829/ The individual VCF files for Sniffles across the samples that are publicly available (not dbGaP) can be found here https://doi.org/10.5281/zenodo.8144524

## Code availability

Source code for Sniffles2 is available at https://github.com/fritzsedlazeck/Sniffles and https://doi.org/10.5281/zenodo.8121996 the auxiliary scripts are available at https://github.com/smolkmo/Sniffles2-Supplement and https://doi.org/10.5281/zenodo.8122060

## Acknowledgments

This research was supported in part by the National Institute of General Medical Sciences (R01 GM132589) to CMBC & FJS. This research was supported in part by the Intramural Research Program of the National Institutes of Health (National Institute of Neurological Disorders and Stroke; project number: 1ZIANS003154). This research was supported in part by the MSA Trust to CP. FJS, MS, MM LP are partially supported by NIH grants (UM1HG008898, 1U01HG011758-01, U01 AG058589, 1UG3NS132105-01 and PO 75N95021P00215) This work was supported in part by the Center for Alzheimer’s and Related Dementias (CARD) within the Intramural Research Programs of the National Institute on Aging (NIA) and the National Institute of Neurological Disorders and Stroke (NINDS), parts of the National Institutes of Health within the Department of Health and Human Services (project ZO1 AG000538-02). This research was funded in part by Aligning Science Across Parkinson’s [Grant number 000430] through the Michael J. Fox Foundation for Parkinson’s Research (MJFF). For the purpose of open access, the author has applied a CC BY [replace with CC0, CC BY-ND if appropriate] public copyright license to all Author Accepted Manuscripts arising from this submission.

## Contributions

MS implemented the software. MS, FJS designed the study. CMBC, CP, MG, DP, SS & FJS generated the data. MS performed the benchmark analysis and LFP performed the data analysis for the population, mosaic and cancer sections. EKE,CP, DH performed PCR validations. MS, LFP, CMG, SB, KH, MM, FJS contributed to data interpretation. All the authors reviewed and edited the manuscript.

## Competing interests

FJS receives research support from PacBio and Oxford Nanopore. S.W.S. is a member of the Scientific Advisory Council of the Lewy Body Dementia Association and an editorial board member of JAMA Neurology, Journal of Parkinson’s Disease, and the Neurodegeneration Specialty Section for Frontiers of Neurology, Frontiers of Psychiatry, and Frontiers of Neuroscience. LFP is sponsored by Genentech, Inc. KH is an employee of Bionano Genomics.

## Notes

### Summary of Updates

We have extensively updated the section of the MSA brain analysis and clarified some of the benchmarking (Figure 1) better. We also added a cancer comparison section (COL829)

